# Profiling Genetic Diversity Reveals the Molecular Basis for Balancing Function with Misfolding in Alpha-1 Antitrypsin

**DOI:** 10.1101/2022.03.04.483066

**Authors:** Chao Wang, Pei Zhao, Shuhong Sun, Xi Wang, William E. Balch

## Abstract

Genetic variation of alpha-1 antitrypsin (AAT) is responsible for alpha-1-antitrypsin deficiency (AATD) leading to gain-of-toxic aggregation in the liver and loss-of-function on <u>n</u>eutrophil <u>e</u>lastase (NE) inhibitory activity in the lung contributing to <u>c</u>hronic <u>o</u>bstructive <u>p</u>ulmonary <u>d</u>isease (COPD) during aging. To probe the molecular basis for how biology designs the protein fold to achieve balance between sequence, function and structure contributing to AATD in the population, we measured the intracellular monomer and polymer, secreted monomer and polymer and NE inhibitory activity of 75 alpha-1-antitrypsin (AAT) variants. To address the complex folding dynamics affecting the form and function of the protein fold that is differentially impacted by variants in the population, we applied a <u>G</u>aussian <u>p</u>rocess <u>r</u>egression (GPR) based machine learning approach termed <u>v</u>ariation <u>s</u>patial <u>p</u>rofiling (VSP). By using a sparse collection of extant variants to link genotype to phenotype, VSP maps <u>s</u>patial <u>c</u>o<u>v</u>ariance (SCV) relationships that quantitate the functional value of every residue in the wild-type (WT) AAT sequence with defined uncertainty in the context of its protein fold design. The SCV-based uncertainty allows us to pinpoint critical short- and long-range residue interactions involving 3 regions-the N-terminal (N1), middle (M2) and carboxyl-terminal (C3) of AAT polypeptide sequence that differentially contribute to the balance between function and misfolding of AAT, thus providing an unanticipated platform for precision therapeutic development for liver and lung disease. By understanding mechanistically the complex fold design of the metastable WT AAT fold, we posit that GPR-based SCV provides a foundation for understanding the evolutionary design of the fold from the ensemble of structures found in the population driving biology for precision management of AATD in the individual.

---

Genetic variation in the human population provides a reservoir for the protein fold to evolve conformational plasticity required for form and function in complex biological environments, but also poises the risk for protein misfolding and/or aggregation that could lead to human genetic disease^1, 2^. Understanding the molecular basis balancing protein function and the propensity for misfolding/aggregation in response to genetic variation in the individual is crucial for precision management of protein misfolding disease^1, 3^.

<u>A</u>lpha-1-<u>a</u>ntitrypsin <u>d</u>eficiency (AATD) is an inherited autosomal recessive disorder driven by polymorphisms in the *SERPINA1* gene that is the principle modifier of chronic obstructive pulmonary disease (COPD)^4^. >600 genetic variants in *SERPINA1* gene are reported in the human population that differentially impact the conformation and/or function of its protein product, <u>a</u>lpha-1-<u>a</u>ntitrypsin (AAT)^3, 5^. AAT is an N-linked glycosylated protein mainly synthesized and folded in the endoplasmic reticulum (ER), the first step of the exocytic pathway, of hepatocytes that comprise ∼80% of the liver cells. <u>W</u>ild-type (WT) AAT (also referred to as the M allele) is secreted into circulation to maintain a plasma concentration of 1-2 g/L^6^. The major function of AAT is that of an antiprotease to <u>n</u>eutrophil <u>e</u>lastase (NE) to prevent tissue degradation in the lung. The most common allele leading to severe forms of AATD is Z variant (E366K). It presents in 1 of 25 individuals with European descent in which 1 of ∼2000 are homozygous leading to severe disease^7^. The Z variant (referred to as Z-AAT) leads to misfolding and intracellular polymerization of AAT in the ER, which can trigger liver disease phenotypes including chronic hepatitis, cirrhosis and, more rarely, hepatocellular carcinoma^8–10^. Due to degradation of misfolded Z-AAT^11, 12^,and/or the accumulation of ordered polymeric inclusions in the ER^13^, only 10 to 15% of Z-AAT in homozygous patients is secreted into the circulation. A deficiency of functional AAT disrupts the protease-antiprotease balance in the lung, triggering emphysema and COPD^14^. Polymers of AAT are also found to be secreted from liver and lung cells^15–21^ that potentially contribute to inflammation and lung damage^22–24^. Compared with efforts focused on Z-AAT, the impact of the many other variants are poorly studied or completely unknown^3, 5, 7^.

As a metastable protein, the different folding conformations of AAT are tightly managed by multiple proteostasis pathways^3, 6, 25^. For example, the calnexin cycle has been shown to be involved in the folding of AAT during nascent synthesis in the ER in response to its N-linked oligosaccharide^25, 26^ and subject to glycosidase modifiers such as ER mannosidase I (ERManI) that accelerates degradation of misfolded AAT^11, 27–30^. Other pathways include the abundant Hsp70 (BiP) and Hsp90 (GRP94) chaperones and their co-chaperones that are highly enriched in the ER^31–33^. While the soluble misfolded Z-variant AAT is degraded by the proteasome through the ER associated degradation (ERAD) pathway^11, 27–30^, intracellular polymer is thought to be disposed of by autophagic pathways^34–36^.

Understanding the differential folding properties of AAT and its many variants in the individual will be essential to tailor the management of its folding pathways for precision development of therapeutics that effectively manage stability, misfolding and/or function of AAT variants^3^. While small molecules have been developed to directly bind to Z-AAT that block polymerization and lead to increased secretion^37–39^, an observation consistent with the ‘loop-sheet’ insertion model for aggregation in the ER^40^, it also abolishes the NE inhibitory activity of AAT^38^. These results indicate the importance of a deeper understanding of the sequence, folding, function and structure relationships spanning the entire protein sequence. An improved appreciation of how nature uses genetic variation over billions of years to evolutionarily balance folding and misfolding to create highly dynamic sequence-to-function-to-structure relationships could reveal new therapeutic strategies to shift the balance away from AAT intracellular polymerization to improve monomer secretion and function providing novel corrective interventional strategies for both liver and lung clinical features responsible for AATD.

To probe the molecular basis for the balance between misfolding and function of AAT, we measured the intracellular monomer and polymer levels, secreted monomer and polymer levels, and NE inhibitory activity for 75 AAT variants distributed across the protein sequence that include both pathogenic and benign variants found in the population. To address the complex folding dynamics likely differentially impacted by these variants, we applied a <u>G</u>aussian <u>p</u>rocess <u>r</u>egression (GPR) based machine learning approach termed <u>v</u>ariation <u>s</u>patial <u>p</u>rofiling (VSP)^1, 41–44^.

By using a sparse collection of extant variants as input to link genotype to phenotype, VSP maps <u>s</u>patial <u>c</u>o<u>v</u>ariance (SCV) relationships that quantitate with assigned uncertainty the functional value of every residue in the context of its global protein fold design. Using the characterized variants as known inputs, SCV allow us to build ‘phenotype landscapes’ that map sequence-to-function-to-structure relationships on a residue-by-residue basis to understand the highly evolved design of the WT protein fold for all uncharacterized residues in the WT sequence. SCV relationships pinpoint critical unknown residue interactions driving AAT folding/misfolding, monomer/polymer balance, secretion and function of AAT. By understanding mechanistically the complex fold design of native AAT using GPR based SCV relationships, we posit that knowledge of the differential contribution of every residue to the diverse folding and functional features contributed by the ensemble of AAT folds found in the population provides a foundation for precision management of AATD in the individual.

## Results

### Assaying the diverse folding and functional features of AAT variants

To generate a collection of AAT variants associated with known clinical features, we utilize 44 pathologic variants associated with AATD liver and lung phenotypes reported in the literature^3, 5, 45^, and those currently annotated in the ClinVar database^46^. We also include 31 variants that are annotated as benign or of ‘uncertain significance’ in the ClinVar database^46^. A total of 75 AAT variants are missense variants with the exception of 3 variants that generate a truncated protein (Y62*, E281* and Null Hong Kong (NHK)) that severely disrupt function (**Fig. 1A**). The collection of variants includes the most common missense variants in AAT in terms of allele frequency (AF) that have been reported in the general population (gnomAD database^47^) including M1 (V237A; 22% AF), M2 (R125H; 15.6% AF), M3 (E400D; 27% AF), S (E288V; 2.3% AF), Z (E366K; 1.1% AF) and others that have >0.1% AF. The residues impacted by the variant collection are spread across the entire AAT sequence (**Fig. 1A**; **Fig. 1B**, brown balls), thus providing molecular fiduciary markers^1^ that enables us to probe the sequence based folding and function space defining the ensemble of AAT structures found in the extant population in response to genetic variation.

**Figure 1.**
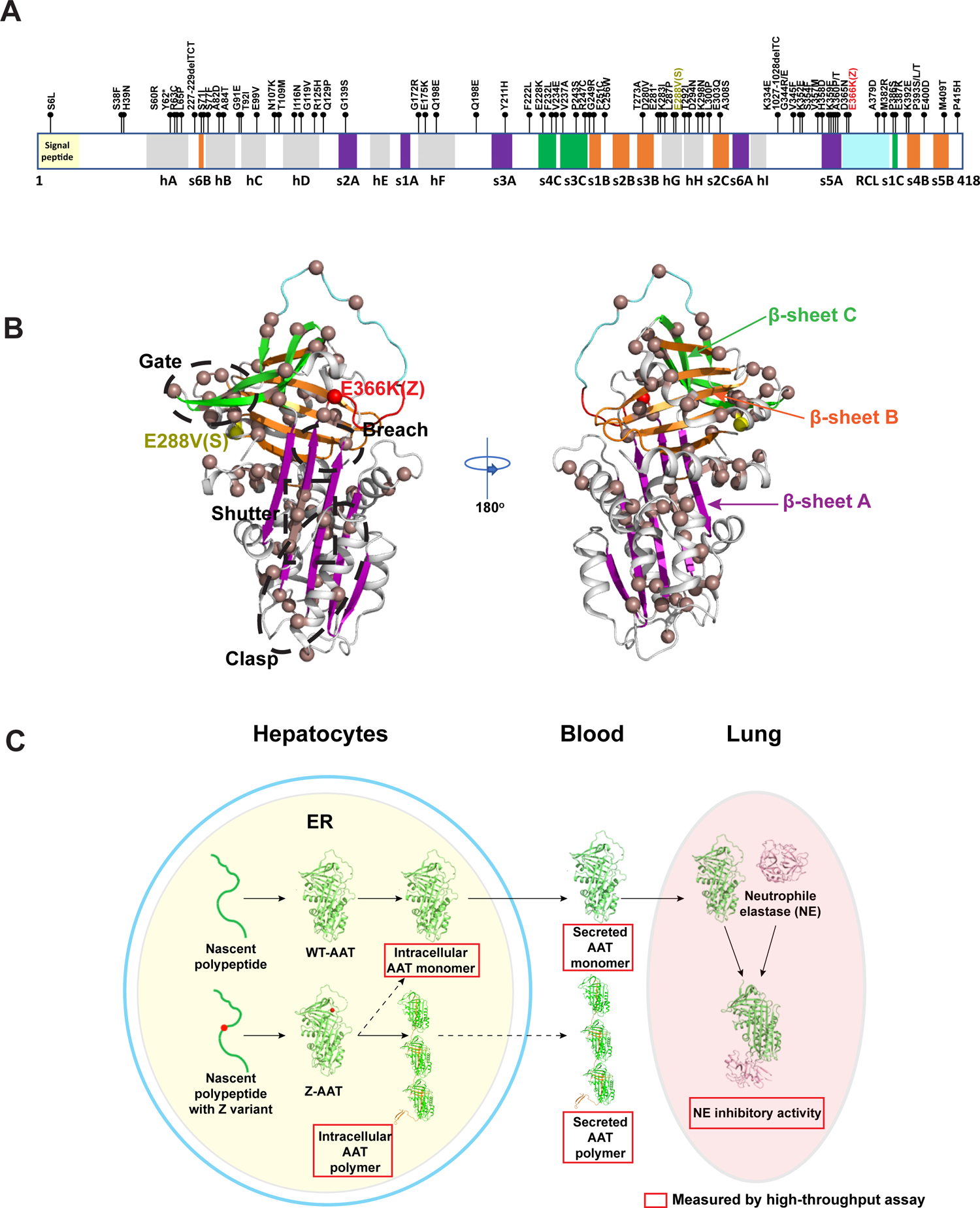
Genetic diversity in AATD. (**A**) 75 AAT variants from the population investigated in this study are distributed at different structural elements across the entire AAT polypeptide sequence. Eight α-<u>h</u>elices (*hA-hI*) are indicated by gray blocks. Three β-<u>s</u>heets comprising *s1-6A (sheet A)*, *s1-6B (sheet B)* and *s1-4C (sheet C*) are highlighted by purple, orange and green respectively. The <u>r</u>eaction <u>c</u>entral loop (RCL) is illustrated by the cyan block. Z (E366K) and S (E288V) variants are labeled in red and yellow respectively. (**B**) Distribution of the variants in the 3D structure of AAT (PDB: 3NE4^70^). The alpha carbon of the variant residues are shown as brown balls. The gate, breach, shutter and clasp regions^72^ are indicated. β-sheets A, B, and C are highlighted. (**C**) WT AAT is synthesized in the endoplasmic reticulum (ER) of hepatocytes and secreted as a monomer into circulation for delivery to lung to perform its NE elastase inhibitory activity. AAT variants such as Z (E366K) lead to intracellular polymerization that reduces AAT secretion and function. We have developed high-throughput assays (indicated by red squares) to measure the intracellular monomer and polymer, secreted monomer and polymer, and NE inhibitory activity for each of the AAT variants.

To characterize the impact of a variant on the status of the AAT protein fold responsible for its monomeric or polymeric states based on its folding, stability and trafficking itinerary through the secretory pathway (**Fig. 1C**), we utilized the monomer specific antibody 16f8 (**Fig. S1A**) and the polymer specific antibody 2C1^48^. These conformation dependent antibodies enabled us to develop an enzyme-linked immunosorbent assay (ELISA) (**Fig. S1A-B**) to measure the level of AAT monomer or polymer in intracellular and extracellular environments (**Fig. 1C**) using high-throughput formats (see **Methods**). To measure the inhibitory activity of secreted AAT variants to NE, its natural substrate in the lung, we used a sensitive fluorogenic NE substrate (Z-AAAA)2Rh110^49^ (**Fig. S1B**) (see **Methods**). A high level of NE inhibitory activity prevents the digestion of the fluorogenic NE substrate resulting in a low fluorescence signal (**Fig. S1C**). Combined, use of the conformation specific ELISA and NE inhibitory activity assays allow us to assess the different folding and functional states of AAT variants across its entire biological itinerary from the ER in the liver to secretion out of the cell for delivery to downstream tissues (**Fig. 1C**).

### Defining the folding and functional properties of AAT variants

To generate a comprehensive understanding of the functional phenotypic properties of AAT in response to its physiologic itinerary (**Fig. 1C**), WT and 75 variants were transfected in either a hepatocyte derived cell line Huh 7.5 lacking endogenous AAT^50–52^, or a human bronchial epithelial lung cell line IB3 that has no detectable endogenous AAT^53^ to capture epithelial pathways potentially contributing to lung function. The level of intracellular or extracellular AAT variants in either monomer or polymer form, and the NE inhibitory activity of the extracellular (secreted) pool of AAT variants were measured (**Fig. 2A, Fig. S2**). The measured activity values for secreted monomer, intracellular polymer and NE inhibitory activity are highly correlated between the liver cell line Huh 7.5 and the lung cell line IB3 (**Fig. S3A**). These results demonstrate that the basic folding and functional features of AAT variants are conserved across different cellular environments. Furthermore, for each variant the intracellular level of monomer and polymer conformations are highly correlated with extracellular monomer and polymer, respectively, in both cell lines (**Fig. S3B**). These results demonstrate that extracellular monomer and polymer conformations of AAT are secreted from each cell type with similar efficacies reflecting the conserved operation of the exocytic pathway. Given the similarity of the conformational features associated with each AAT variant that differentially influences the state of its fold design and function, we use the measured levels of intracellular polymer, secreted monomer and the NE inhibitory activity from Huh7.5 cells henceforth to understand how, on a residue-by-residue basis, the AAT fold design is coupled to function to drive different disease states found in the AATD population.

**Figure 2.**
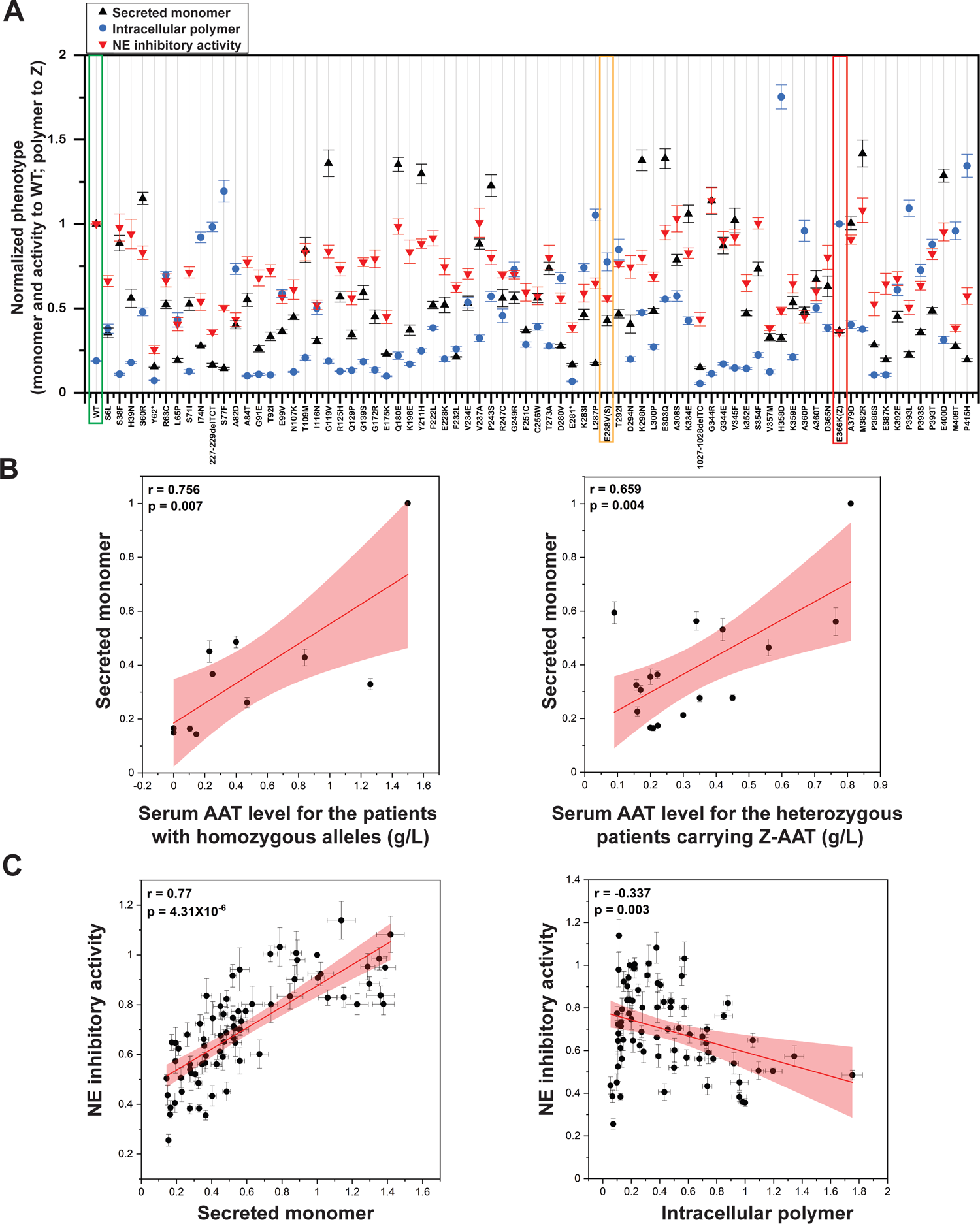
Quantification of secreted monomer, intracellular polymer and NE inhibitory activity for AAT variants. (**A**) The levels of secreted monomer, intracellular polymer and NE inhibitory activity for WT-AAT and 75 AAT variants transfected in Huh7.5 cells are shown (see **Methods**). The secreted monomer and NE inhibitory activity are normalized to WT values. The intracellular polymer is normalized to Z-AAT value. (**B**) Correlation between measured secreted monomer levels and reported serum AAT levels in AATD patients who are homozygous with the indicated variant genotype (left panel), or from heterozygous patients who share the common Z allele (right panel). The Pearson’s r value and the corresponding p-value are indicated. (**C**) Correlation between the NE inhibitory activity of AAT variants and the secreted monomer (left panel) or intracellular polymer levels (right panel). The Pearson’s r value and the corresponding p-value are indicated.

We first compared WT-AAT to Z-AAT. As expected, Z-AAT shows low monomer secretion (∼37% of WT) and low NE inhibitory activity (∼36% of WT), while WT-AAT shows low intracellular polymer (∼20%) when compared with Z-AAT (**Fig. 2A**). These results are consistent with the severe liver and lung disease phenotypes for patients with homozygous Z triggered by the combined gain of toxicity in the liver due to accumulation of Z-AAT intracellular polymer and loss of function in the lung due to Z-AAT deficiency^3, 7^. In contrast, the S allele (E288V) shows a median level of secreted monomer, intracellular polymer and NE inhibitory activity (**Fig. 2A**), consistent with the mild disease phenotypes of patients with homozygous S allele. We found that our cell-based measurements of secreted monomer levels are significantly correlated with reported AAT serum levels from patients with different variants, indicating the utility of our cell-based assay to capture the clinical features of disease in the patient population (**Fig. 2B**). Moreover, the secreted monomer levels strongly correlate with their NE inhibitory activity (**Fig. 2C**, left panel, Pearson’s r = 0.77, p = 4.3 x 10^-6^), indicating that the monomer conformation we measure corresponds to the functional form. In contrast, the intracellular polymer levels of the variants are moderately anti-correlated with the extracellular NE inhibitory activity values (**Fig. 2C**, right panel, Pearson’s r = −0.34). While variants with high intracellular polymer (**Fig. 2C**, right panel, polymer > 1) generally show low extracellular NE inhibitory activity (< 60% of WT), variants with low intracellular polymer (**Fig. 2C**, right panel, polymer < 0.2) have very diverse NE inhibitory activities ranging from ∼26% of WT to 114% of WT (**Fig. 2C**, right panel). These results suggest that a low polymerization propensity does not necessarily confer high functional activity, suggesting a more complicated set of relationships between folding, stability and the functional properties of AAT variants that contribute to changes in the protein fold design responsible for disease in the AATD population (**Fig. 2C**, right panel).

### Understanding function of each residue spanning the entire AAT sequence using VSP

The distinctive functional properties imposed by natural AAT variants (**Fig. 2**) suggest the potential for differential roles for uncharacterized residues in the sequence contributing to the global ensemble of AAT structures impacting function found in the population. In order to understand the role of all residues shaping WT protein fold design and how they talk to one another to generate a functional structure, we applied variation spatial profiling (VSP)^1^. VSP is a <u>G</u>aussian <u>p</u>rocess <u>r</u>egression (GPR)-based machine learning approach that uses a sparse collection of natural variants and their phenotypes that disrupt normal AAT sequence, function and structure features to generate spatial covariance (SCV) relationships that allows us to build high resolution ‘phenotype landscapes’ on a residue-by-residue basis with defined uncertainty for the entire sequence^1^. The phenotype landscape generated through a GPR principled computational strategy uses the known sparse collection of characterized variants to assign function to the remaining uncharacterized residues in the WT AAT sequence to understand their collective behavior in programming the fold for function in health and disease.

Given the potential impact of variation on the generation of secreted monomer and its level of NE inhibitory activity essential for normal lung function, we first used the phenotypes of each of the 75 variants expressed in Huh7.5 cells (**Fig. 2A**) to generate a 2-dimensional (2D) view of GPR-based phenotype landscape that profiles the SCV relationships between secreted monomer (**Fig. 3A**, *y*-axis) and NE inhibitory activity (**Fig. 3A**, z-axis presented as a color scale) for each residue in the context of the full-length polypeptide sequence calibrated to a scale of 1 (N- to C-terminus including the signal peptide) (**Fig. 3A**, *x-axis*, <u>var</u>iant <u>seq</u>uence <u>p</u>osition (VarSeqP)). The phenotype landscape encompassing the entire AAT polypeptide sequence reveals that AAT variants with low secreted monomer generally have low NE inhibitory activity across the entire polypeptide sequence (**Fig. 3A**, yellow-orange-red), consistent with the significant linear correlation between these features based on only a sparse collection of variants used as the training set (**Fig. 2C**). Here, GPR not only generates the phenotype prediction for understanding mechanistically known SCV based residue interactions, but also assesses the prediction variance for functional relationships for all unknown interactions between residues in the AAT sequence with assigned uncertainty (confidence) (**Fig. 3A**; Pearson’s r = 0.758, p=7×10^-15^, leave-one-out cross validation (LOOCV)) (see **Methods**). The prediction variance value depends on the clustering effect of the input variants, where more clustered variants yield lower variance (higher confidence) for the prediction. The prediction variance values for the top 25% confidence predictions are plotted using contour lines in the landscape (**Fig. 3A**). These high confidence relationships reveal three major regions found at the N-terminal, middle and C-terminal regions of the polypeptide sequence (**Fig. 3A**, numbered brackets) where clustered sequence relationships generally confer low secreted monomer (**Fig. 3A**, y < 0.6) and deficient NE inhibitory activity (**Fig. 3A**, orange-red). These results suggest that there are specific regions along the polypeptide sequence critical for coupling monomer secretion to NE inhibitory activity of AAT that are disrupted by natural variants in the population that cooperate with uncharacterized residues dictating the global function of the AAT fold.

**Figure 3.**
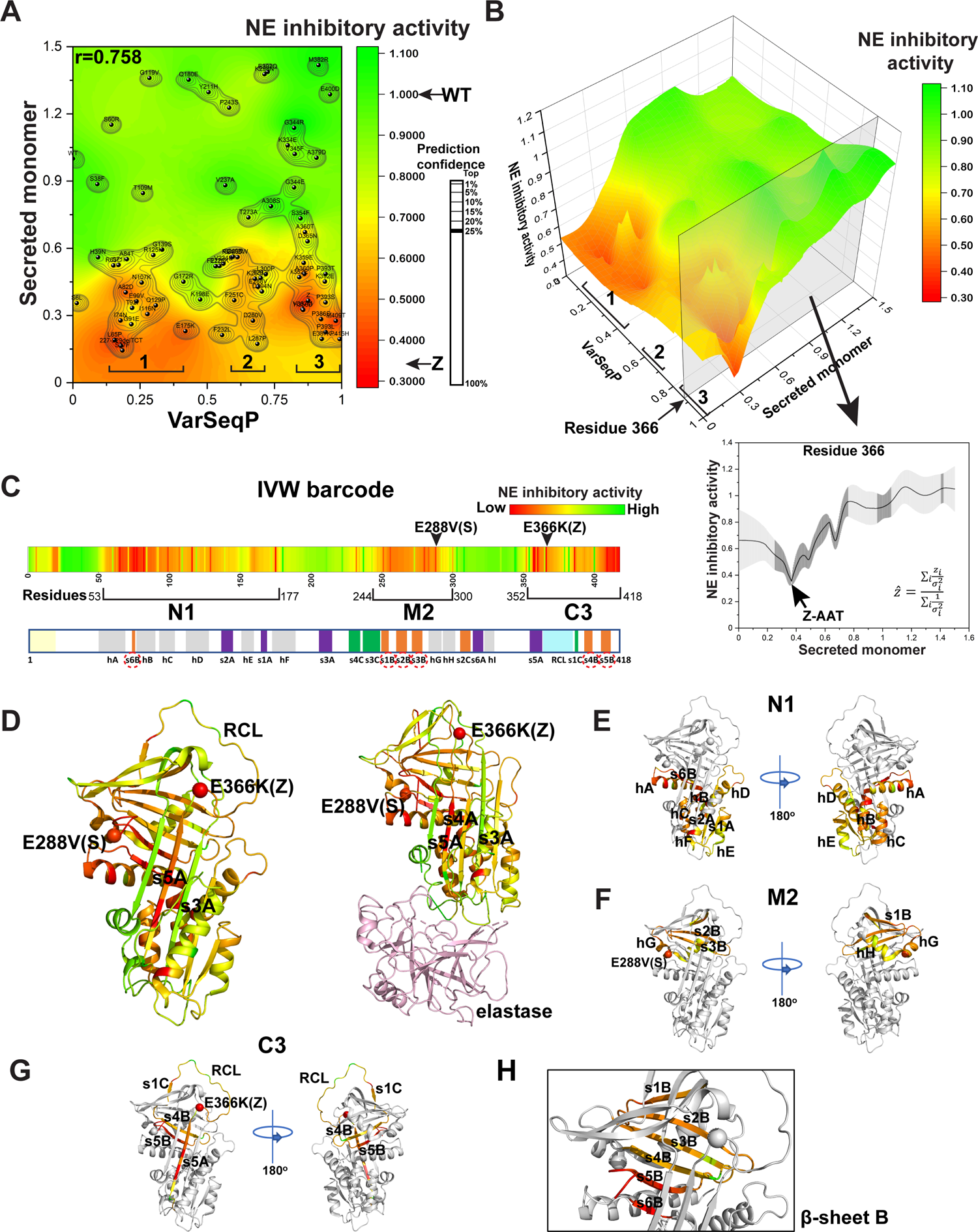
SCV contribution of each residue in the AAT polypeptide sequence to function using GPR. (**A**) A phenotype landscape was built based on input of 75 AAT variants (Fig. 1) by linking the secreted monomer (*y*-axis) to NE inhibitory activity (color scale, z-axis) relative to the <u>var</u>iant <u>seq</u>uence <u>p</u>osition (VarSeqP) (*x*-axis) using GPR-based VSP^1^ (see **Methods**). The VarSeqP is normalized to full-length polypeptide sequence that is assigned a value of ‘1’. The GPR-generated variance for each prediction within top 25% confidence is illustrated by contour lines. Three sequence regions with clustered variants that have low secreted monomer and activity are labeled as ‘1’, ‘2’ and ‘3’. (**B**) The 2D landscape in (A) can be rendered as a 3D landscape by visualizing the *z*-axis reflecting the NE inhibitory activity (color gradient). The gray slice presents the full range of secreted monomer and NE inhibitory activity for residue 366 based on GPR generated predictions from variants used in this study. These values can be plotted as a curve (inset) with the GPR prediction variance shown as error bars (see **Methods**). The prediction variance values within the top 25% confidence contour are highlighted as dark gray (inset). The value of Z-AAT is indicated. The equation used for inverse variance weighting (IVW) is indicated (inset) (see **Methods**). (**C**) The IVW averaged NE inhibitory activity for each residue from the N-terminal to the C-terminal of AAT polypeptide is shown as an ‘IVW barcode’ with the color scale from red to green representing NE inhibitory activity ranging from low (Z-variant AAT) to high (WT AAT). The regions containing residues that are < 75% WT NE inhibitory activity found in the N-terminal, middle, and C-terminal of the AAT polypeptide are labeled as N1, M2 and C3 respectively. **(D-H)** The IVW averaged NE inhibitory activity values for each residue are mapped on the 3D structure of AAT (**D**, left panel; PDB:3NE4^70^) and the AAT-NE complex (**D**, right panel; PDB:2D26^71^) with the structural regions of N1 (**E**), M2 (**F**), C3 (**G**) and β-sheet B (**H**) highlighted.

The 2D phenotype landscape can be rendered as a 3-dimensional (3D) projection to quantitatively visualize the range of predicted SCV relationships of NE inhibitory activity (**Fig. 3B**, *z*-axis) relative to the levels of secreted monomer (**Fig. 3B**, *y*-axis) generated by GPR for every residue in the AAT sequence (**Fig. 3B**, *x*-axis). This provides an ensemble view of function relationships based on SCV relationships defining residue variation found across the population. For example, we can visualize the predicted monomer and activity relationships with probabilities for residue 366 as a curve (**Fig. 3B**, gray slice panel and inset). The error bars represent GPR prediction variance, and the top 25% confidence values are highlighted as dark gray (**Fig. 3B**, inset, see **Methods**). These high confidence regions describing as a collective functional SCV relationships generated by the known input variant in each region, such as Z-AAT with low monomer and activity levels, and the GPR based SCV impact from surrounding input variants reveals the ensemble of monomer and activity variance at the residue 366 position responding to global fold design (**Fig. 3B**, inset).

The GPR generated prediction variance associated with every residue for the full phenotype space enables us to compute an averaged predicted NE inhibitory activity for each residue through inverse variance weighting (IVW) (**Fig. 3B**, inset) (see **Methods**). Using the predictions within the top 25% confidence region (**Fig. 3B**; inset, dark gray regions), IVW prioritizes the assigned value for each residue with low GPR prediction variance (high confidence) given higher value over those with a high GPR prediction variance (low confidence) as illustrated for residue 366 (**Fig. 3B**, inset) to obtain an averaged NE inhibitory activity across the ensemble. We apply the IVW approach to every residue, and the IVW averaged NE inhibitory activity for each residue is mapped to the full-length polypeptide (**Fig. 3C**) as a linear ‘IVW barcode’ map spanning the entire polypeptide sequence (**Fig. 3C**, IVW barcode). The IVW barcode map quantitatively defines 3 high confidence clusters (**Fig. 3C**; labeled brackets, N1, M2 and C3) captured first by SCV relationships in the phenotype landscape (**Fig. 3A**, numbered brackets) that contributes to NE inhibitory activity. As N1, M2 and C3 are located in the N-terminal, middle, and C-terminal regions, respectively, these results suggests that AAT has evolved both short and long range cooperative interactions to manage NE inhibitory activity. While short-range SCV relationships likely emphasize the function of domains that participate in local folding dynamics, the long-range activities connecting the different clusters reflect the complex fold design necessary to integrate nascent synthesis, folding and trafficking in the liver to downstream function in serum and the lung.

### N1, M2 and C3 cooperate in the function-based structural design of AAT

To address how the N1, M2 and C3 regions cooperate from a structural point-of-view with each other to manage AAT protein folding of the nascent chain in the ER with function in downstream environments such as serum and lung, we mapped the IVW averaged values defining AAT NE inhibitory activity for each residue (**Fig. 3C**, IVW barcode) onto the AAT structure (**Fig. 3D**) in the absence (**Fig. 3D**, left panel) or presence (**Fig. 3D**, right panel) of NE. The N1 region consists of the first five *α*-helices, *hA-E*, and two β-strands, *s1A* and *s6B*, of AAT (**Fig. 3E**). The M2 region is composed of three β-strands in β-sheet B including *s1B*, *s2B* and *s3B* and *α*-helices G and H (*hG-H*) where the prevalent E288V S variant is located that renders partial inactivation of AAT function with reduced load of intracellular polymer relative to Z-AAT (**Fig. 3F**). The dominant E366K Z variant allele residue is located in the C3 region that includes in addition to β-<u>s</u>trand 5A (*s5A*) and the reaction central loop (RCL), three β-strands at the C-terminal end (**Fig. 3C**; *s1C, s4B, s5B,* red dotted circles; **Fig. 3G**). After binding of NE and cleavage of the RCL, the cleaved RCL is inserted into β-sheet A forming a new β-strand (*s4A*) that interacts with *s5A* (**Fig. 3D**, right panel), irreversibly capturing the inactivated extracellular NE for disposal by the endocytic-lysosomal pathway^54–56^. GPR-based SCV relationships defined by IVW barcode mapping underscore the high value of the C3 cluster in folding and function, demonstrating the importance of not only the *s5A* and RCL regions, but emphasize the unanticipated importance of the β-strands *s4B* and *s5B* at the C-terminal, suggesting that more complex dynamics occur between the structural elements found in C3 to achieve the NE activity of AAT.

Strikingly, though N1, M2 and C3 are separated in the primary sequence in the N-terminal, middle and C-terminal regions, respectively, of the polypeptide (**Fig. 3C**), they interact with each other through the β-sheet B (**Fig. 3C**, *s6B* strand for N1, *s1-3B* strand for M2 and *s4-5B* strand for C3, red dotted circles; **Fig. 3H**). The integration of each of the core SCV regions through β-sheet B indicates that the protein fold design of AAT is based on elements that contribute independent as well as dependent properties-providing the first glimpse of evolved functional relationships now captured by GPR to understand the spatial design of the functional AAT fold and how variants may impact health and disease progression.

### N1, M2 and C3 facilitate AAT monomer secretion

To understand the relationship between retention of NE inhibitory activity in secretion of monomer on a residue-by-residue basis, we used GPR to understand the role of sequence position of AAT variants (**Fig. 4A**, *x*-axis) using NE activity of each variant as the *y*-axis coordinate (**Fig. 4A**) to define the SCV relationships contributing to monomer secretion as the *z*-axis coordinate (**Fig. 4A**, *z*-axis color scale) to generate a monomer secretion phenotype landscape (**Fig. 4A**). Consistent with the NE inhibitory activity landscape (**Fig. 3A**), the monomer secretion phenotype landscape also revealed three major cluster regions that affect monomer secretion (**Fig. 4A**; N1, M2 and C3) leading to reduced extracellular monomer levels. IVW was used to generate an averaged predicted monomer secretion value for each residue (**Fig. 4B**) and the corresponding structural maps (**Fig. 4C-G**). They show strong functional overlap with NE inhibitory activity (compare **Fig. 4C-G** with **Fig. 3C-H**). It is now clear that both short and long-range effects of N1, M2 and C3 clusters that effect function also contribute to the status of monomer secretion (**Fig. 4C-G**). These results emphasize that monomer secretion and NE inhibitory activity are by genomic sequence design structurally integrated on a comprehensive residue-by-residue basis spanning the entire sequence.

**Figure 4.**
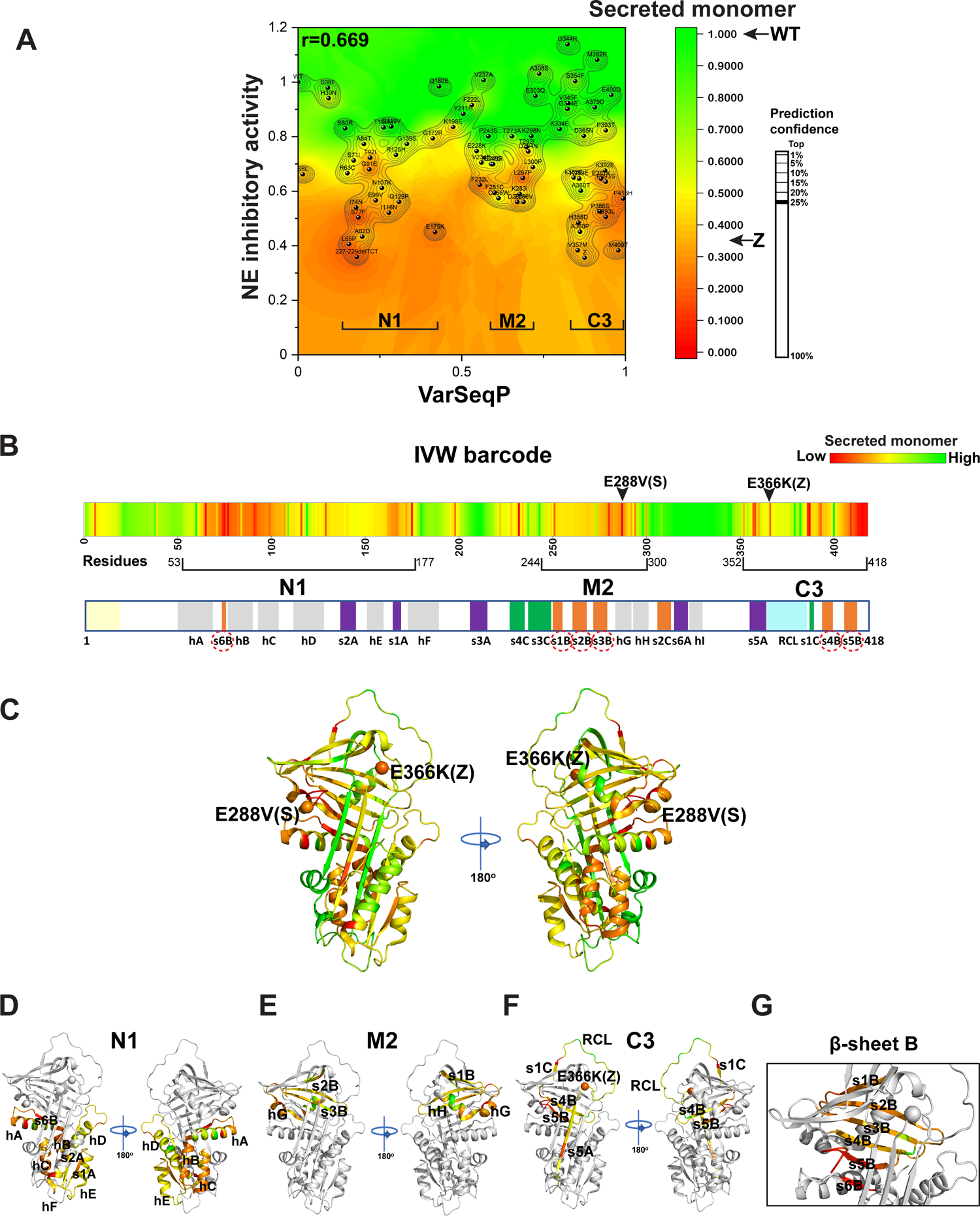
The residue-by-residue SCV contribution to secreted monomer. (**A**) Shown is the phenotype landscape linking NE inhibitory activity (*y*-axis) to secreted monomer (color scale, z-axis) across the entire AAT polypeptide sequence (*x*-axis). The prediction confidence within the top 25% confidence interval is defined by contour lines. (**B**) The IVW averaged secreted monomer levels for each residue are presented from N-terminal to C-terminal of AAT polypeptide as a barcode with the color scale from red to green representing low (Z-variant AAT) to high (WT AAT secretion of monomer. (**C-G**) The IVW averaged monomer secretion levels for each residue are mapped on the 3D structure of AAT (C; PDB:3NE4^70^), with the structural regions N1 (**D**), M2 (**E**), C3 (**F**) and β-sheet B (**G**) highlighted.

### GPR captures the molecular mechanism for AAT intracellular polymerization

Given the short and long range interactions required for AAT monomer secretion and NE inhibitory activity, we examined how these interactions contribute to polymerization of AAT that drives the liver disease clinical phenotype. To understand the molecular basis for AAT intracellular polymerization, we used our sparse collection of variants and GPR to profile the SCV relationships linking secreted monomer (**Fig. 5A**, *y*-axis) with intracellular polymer load (**Fig. 5A**, z-axis color scale) to predict the contribution of every residue spanning the AAT polypeptide (**Fig. 5A**, *x*-axis) contributing to intracellular polymerization. Strikingly, the resulting monomer-polymer phenotype landscape (**Fig. 5A**) is very different from the phenotype landscapes predicting the role of each residue in NE inhibitory activity (**Fig. 3A**) or monomer secretion (**Fig. 4A**). Indeed, a leave-one-out cross-validation (**Fig. 5A**, Pearson’s r = 0.317, p = 0.007) showed only a moderate statistically significant relationship when compared with the strong statistical significance observed for the for NE inhibitory activity phenotype landscape (**Fig. 3A**). These results imply there are different SCV relationships driving intracellular polymer related liver disease from the NE inhibitory activity related lung phenotypes. Using IVW (**Fig. 5B**), the number of residues in the N1 and M2 clusters contributing to intracellular polymerization (**Fig. 5A**, **B**; orange to red) are reduced when compared to the number of residues critical for monomer secretion (**Fig. 4A**, **B**; orange to red) and NE inhibitory activity (**Fig. 3A**, **C**; orange to red). In contrast, the C3 cluster of residues, that contribute substantially to deficient monomer secretion and NE inhibitory activity (**Fig. 3A**, **C**; **Fig. 4A**, **B**), also contribute substantially to the level intracellular polymerization including the β-strands *s4B* and *s5b* (**Fig. 5B**, dotted red circles).

**Figure 5.**
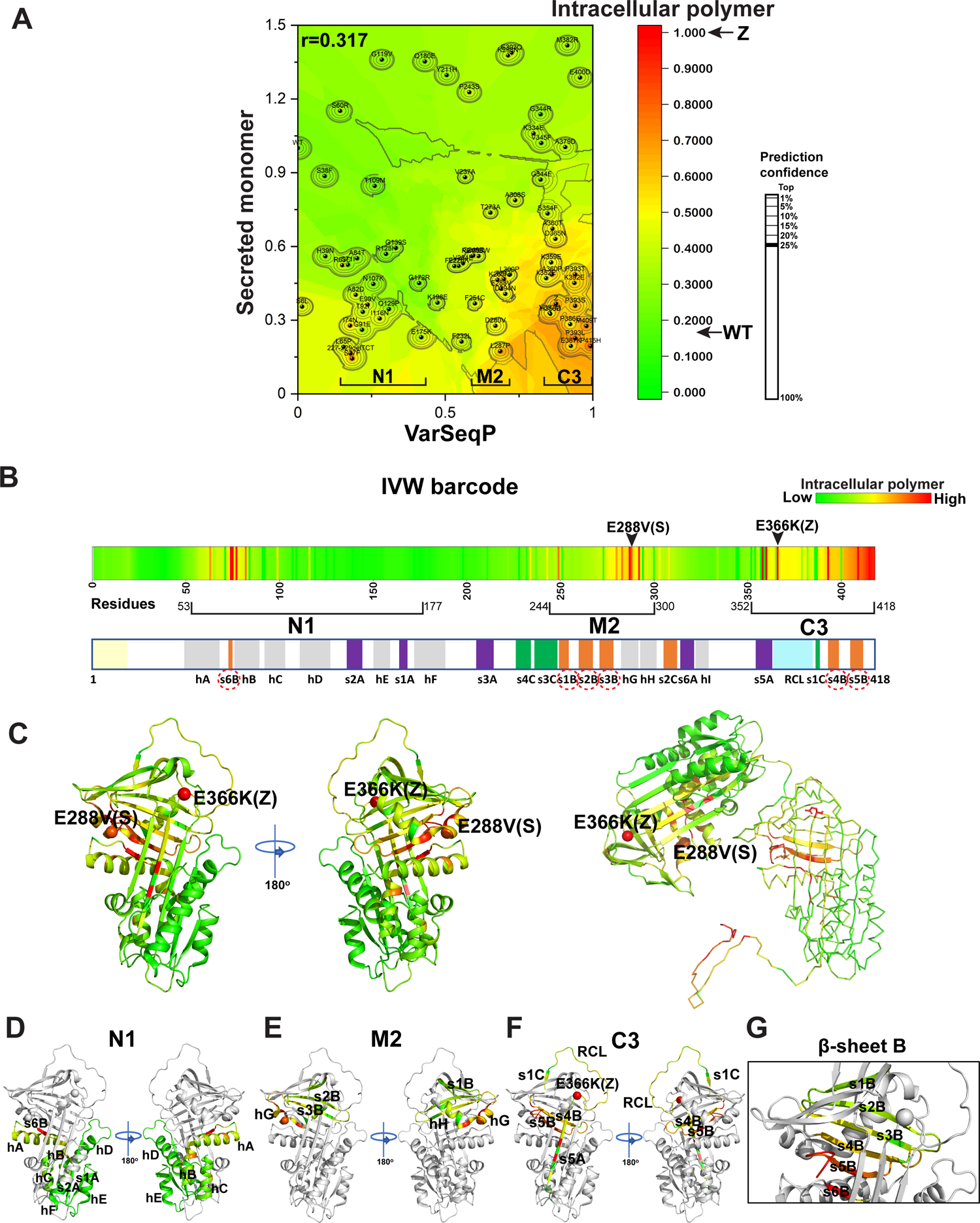
Residue-by-residue SCV relationships reveal the intracellular polymerization mechanism. (**A**) Phenotype landscape linking secreted monomer (*y*-axis) to intracellular polymer (color scale, z-axis) across the entire AAT polypeptide sequence (*x-*axis). The N1, M2 and C3 regions that were identified by low NE inhibitory activity are indicated in the landscape. The top 25% confidence interval is defined by contour lines. (**B**) The IVW averaged intracellular polymer levels for each residue are presented from N-terminal to C-terminal of the AAT polypeptide as a barcode with the color scale from red to green representing high (Z-variant AAT) to low intracellular (WT AAT) polymer levels. (**C-G**) The IVW averaged intracellular polymer levels for each residue are mapped on the 3D structure of AAT (**C**, left panel; PDB:3NE4^70^) or AAT intracellular polymer (**C**, right panel; PDB:3T1P^57, 58^) with the structural regions N1 (**D**), M2 (**E**), C3 (**F**) and β-sheet B (**G**) highlighted.

To understand structurally the impact of each residue in the AAT sequence leading to intracellular polymerization, the IVW based intracellular polymer score (**Fig. 5B**) for each residue was mapped to the AAT structure (**Fig. 5C-H**). Strikingly, in addition to the RCL and *s5A*, known to be important for the ‘loop-sheet’ insertion NE inhibitory mechanism, the SCV relationships defined by the phenotype landscape highlights a central role for the β-sheet B (**Fig. 1B**, β-sheet B) in the mechanism of AAT intracellular polymerization (**Fig. 5C**). Importantly, the residues critical for intracellular polymer formation in β-sheet B comprises *s6B* from N1, *s3B* from M2 and *s4-5B* from C3 (**Fig. 5B**, dotted red circles**; 5D-G**), suggesting disruption of the long-range interactions in β-sheet B through sequence variation triggers intracellular polymer formation for which C3 contributes a significant role. Moreover, the SCV relationships also suggest that the interactions between the β-sheet B and alpha helices *hG-hH* in the M2 region encompassing the S-allele are important in determining the intracellular polymer state of AAT variants (**Fig. 5C** and **5E**). These observations are consistent of the recent cryo-EM study of the *in vivo* Z-AAT intracellular polymer isolated from human Z-variant hepatocytes^57^, which suggested that unlike the loop-sheet insertion model involving the RCL, the C-terminal two β-strands (*s4B* and *s5B*) insert into the β-sheet B of another molecule in the absence of NE substrate, referred to as the C-terminal model^57, 58^ (**Fig. 5C**, right panel). These results indicate the power of GPR based SCV relationships using only a sparse collection of natural variants to capture the fundamental design features of the AAT metastable fold during nascent synthesis and maturation in the ER likely impacting the intracellular polymeric state of the fold driving progression of disease during aging (**Fig. 5C**).

### Cycloheximide (CHX) chase experiments show regional regulation of conformation and function relationships

To understand whether N1, M2 and C3 regions highlighted by our GPR analyses show the predicted differential conformational properties affecting AAT behavior, we performed CHX chase experiments for selected variants in these regions that have lower monomer secretion and NE inhibitory activity when compared to WT values (**Fig. 6A**; **Fig. 2A**). We quantitated the intracellular monomer level of these variants over the CHX chase time course (**Fig. 6B**) (see **Methods**). These results show that N1 variants (**Fig. 6B**; blue lines) show a faster rate of loss for the intracellular monomer when compared with C3 variants (**Fig. 6B**, red lines) and M2 variants (**Fig. 6B**, yellow lines), suggesting that perturbation of the stability in the N-terminal region of AAT causes rapid degradation of misfolded monomer, perhaps by disrupting the fold during nascent synthesis. In contrast, quantification of the intracellular polymer over the CHX chase time course shows that the C3 variants result in lower rate of loss (**Fig. 6C**, red lines) when compared with the M2 and N1 variants. This result is consistent with an SDS-PAGE analysis of AAT stability based on immunoblotting (**Fig. S4**) where the C3 variants have the slowest rate degradation among variants tested for the immature glycosylated form found in the ER (**Fig. S4B**), for the total intracellular AAT level (**Fig. S4C**), and for the ratio between the immature glycosylated form and total intracellular AAT (**Fig. S4D**). These results suggest that the intracellular polymer formed by C3 variants (that includes the Z variant) can acquire a more stable conformation, highlighting the key role of C3 in driving the co- and/or post-synthesis intracellular polymerization and consequential severe disease.

**Figure 6.**
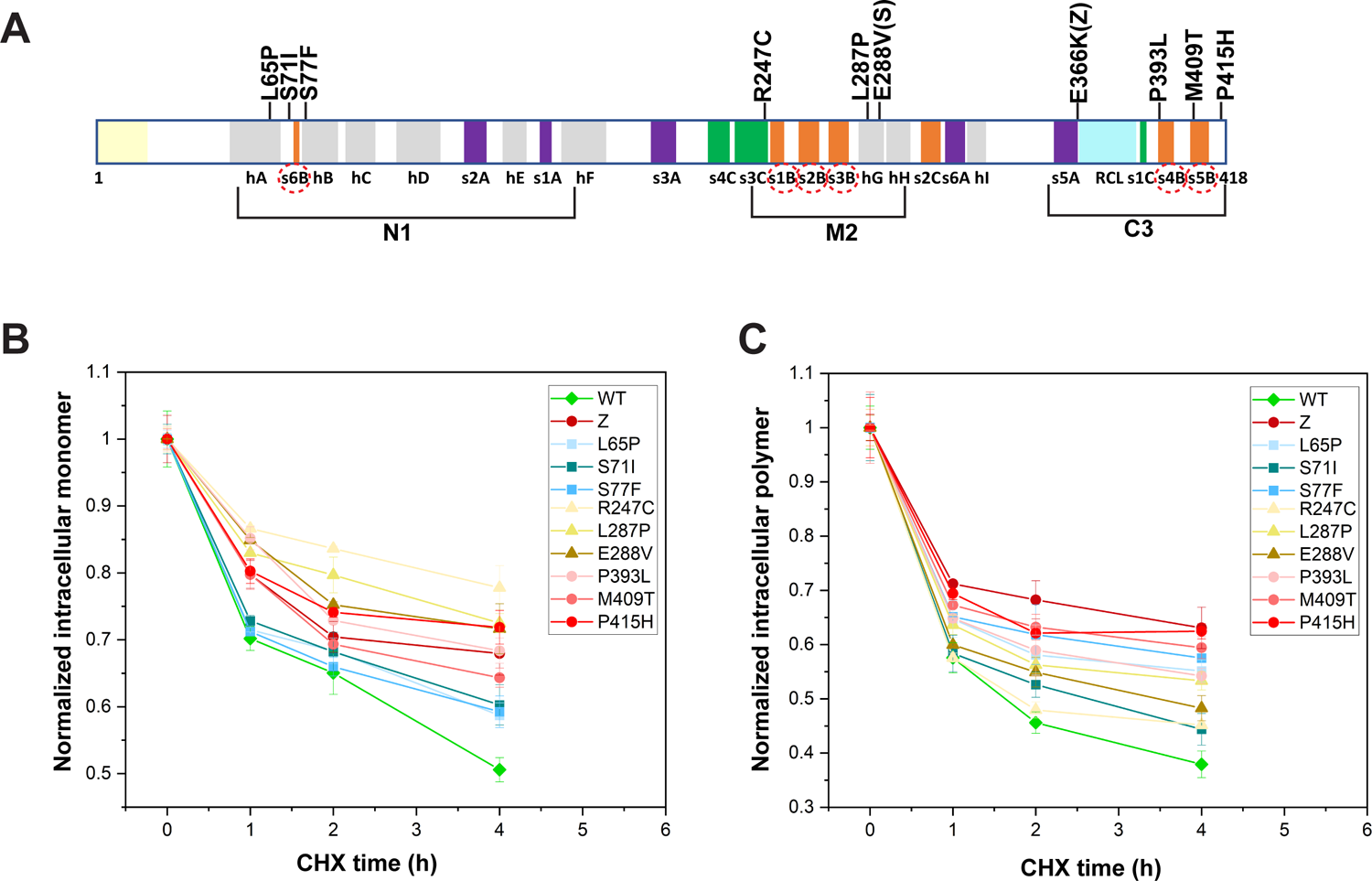
Cycloheximide (CHX) chase reveals the protein stability for select N1, M2 and C3 variants. Select variants from N1, M2 and C3 regions (**A**) were transfected to the Huh7.5 AAT knock-out cell line for 48 h and then treated with CHX (50 µM) for 0, 1, 2 or 4 h. The intracellular sample were collected for ELISA detection of levels of monomer (**B**) and polymer (**C**), which are normalized to the levels prior to CHX treatment (0 h).

### A global map of residues contributing to AAT form versus function in disease

To address more comprehensively the role of different regions on intracellular polymer formation, monomer secretion, and NE inhibitory activity at residue-by-residue resolution, we normalized the GPR-based phenotype value for each residue to the distribution of all the residues by computing a ‘standard score’ (**Fig. 7A**) (see **Methods**). The standard score for a given residue and its associated feature defines the number of standard deviations away from the mean distribution of that feature. The mean corresponds to ∼74% of WT activity and ∼64% WT monomer or ∼40% Z-AAT intracellular polymer, consistent with the fact that our collection of variants comprises both pathogenic and benign variants as representative of the alleles characterizing the AAT variation in the population (**Fig. 7A**). A standard score of ‘+1’ indicates that the GPR-based phenotype value of this residue is one standard deviation away from the mean towards WT phenotype (or at ∼84.1 percentile of all residue phenotypes) (**Fig. 7A**). In contrast, a ‘-1’ standard score indicates one standard deviation away from the mean towards Z-AAT phenotype (or at ∼15.9 percentile of all residue phenotypes) (**Fig. 7A**). By using the standard score, we can directly compare the different folding, secretion, and NE activity features on a common quantitative platform to understand mechanism of action of a particular residue in the ensemble of AAT folds contributing to health and disease across the population.

**Figure 7.**
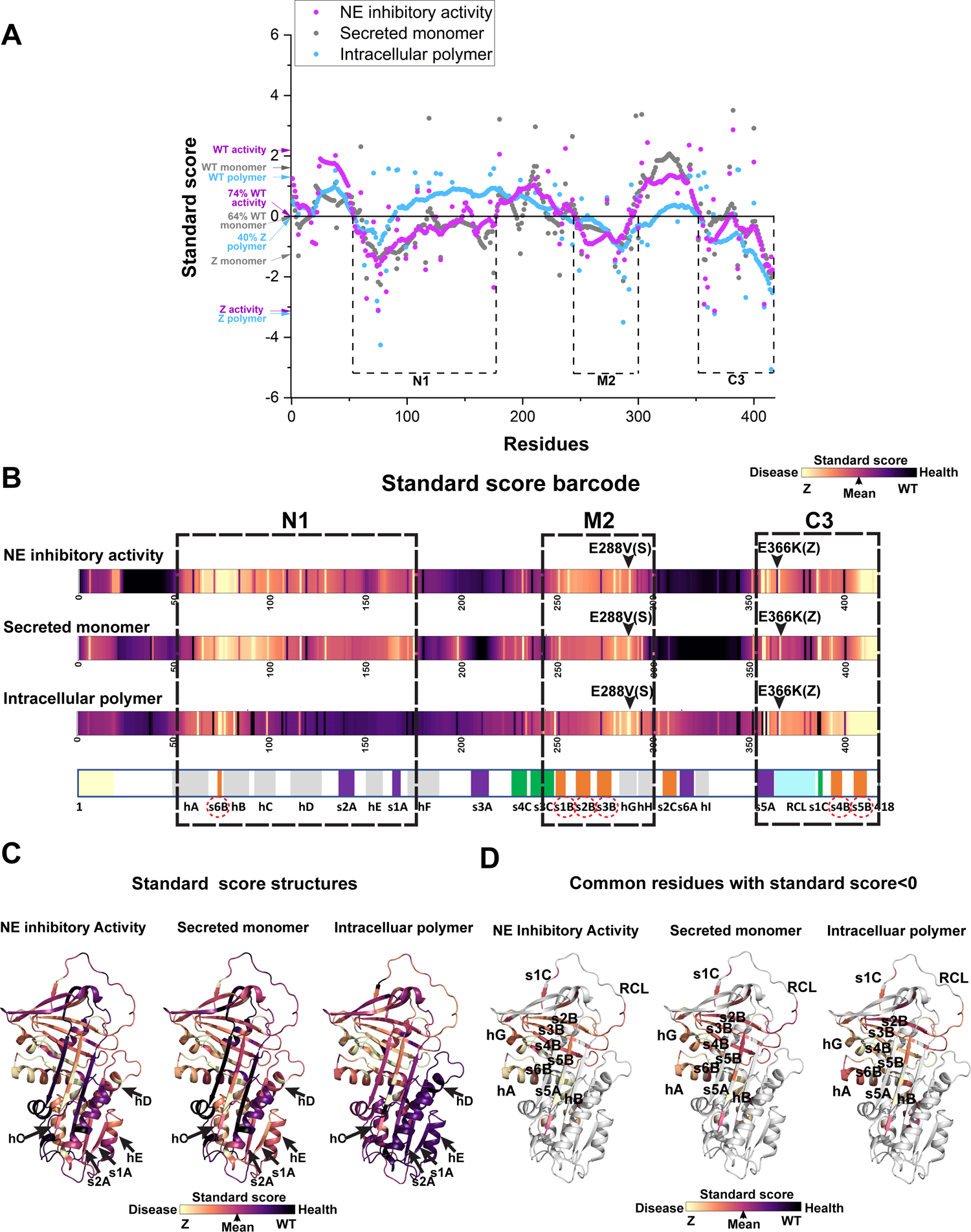
Common and differential residue contributions to AAT phenotypes. (**A**) Standard scores (see **Methods**) define the number of standard deviations by which the IVW averaged phenotype value of each residue is above or below the mean value of the distribution for each phenotype. The standard scores for WT and Z-variant for each phenotype is indicated and the mean value for each phenotype labeled. The regions N1, M2 and C3 that have standard scores of NE inhibitory activity generally below the mean are highlighted by dotted boxes. (**B-C)** The standard scores for each residue are mapped as a barcode (**B**) and on the 3D structure of AAT (**C**). Structural elements in N1 domain in management of the monomer secretion and NE inhibitory activity, but not polymer formation are highlighted by arrows in (**C**). (**D**) Common residues that have standard score below mean (<0) for NE inhibitory activity, monomer secretion, and polymer formation are highlighted.

As expected, the standard scores of NE inhibitory activity for every residue (**Fig. 7A**, purple) generally overlap with those observed for secreted monomer (**Fig. 7A**, grey), consistent with the fact that these two phenotypes are highly correlated (**Fig. 2C**). The residues in N1, M2 and C3 regions generally have standard scores lower than mean (‘0’) for NE inhibitory activity and secreted monomer, consistent with the observation that residue activity in these regions are coordinated to generate the secreted, functional fold (**Fig. 7A**, purple and grey curve). In contrast, the standard score curve for all residues impacting the level of intracellular polymer formation show very distinct trends in specific regions (**Fig. 7A**, light blue curve) that differentiate function from structure. For example, a large sequence region within N1 has the standard score for intracellular polymer formation above 0 with a much higher score than that of secreted monomer and NE inhibitory activity (**Fig. 7A**). These results indicate that the functional deficiency contributed by these residues is not due to generation of intracellular polymer, rather reflects a rapid degradation of misfolded monomer (**Fig. 7A**). Another differential region starts from the end of M2 region (**Fig. 7A**) where the standard scores for monomer secretion and NE inhibitory activity are much higher than that of intracellular polymer, illustrating that there are regions of the AAT structure design that contribute independently to intracellular polymer formation from their roles in secretion and activity.

Using the standard scores (**Fig. 7A**) for each residue we generated a ‘standard score barcode’ map of AAT fold design to illustrate the contribution of different structural elements to different phenotypes (**Fig. 7B**). The C3 region spanning from *s5A*-*RCL* to *s4B-s5B* in the C-terminus of AAT polypeptide has standard scores generally below the mean reflecting a Z-AAT like phenotype across all the measured features. These results suggest it is critical in driving intracellular polymer formation resulting in a deficiency in monomer secretion and NE inhibitory function (**Fig. 7B**). M2 also has shared region with a Z-AAT-like disease phenotype that runs from the end of *s3B* to *hG* where the S allele (E288V) is located (**Fig. 7B**). Interestingly, the residues after *hG* contribute to polymer formation but do not strongly impact the monomer secretion and NE inhibitory activity (**Fig. 7B**). The largest differences in terms of residue contributions to different phenotypes are observed in N1 region (**Fig. 7B**). While β-strand *s6B* and its flanking α-helices *hA* and *hB* show Z-AAT-like disease standard scores for all the measured phenotypes, the residues following *hB* including *hC, hD, s2A, hE, s1A* have a ‘healthy’ standard score for polymer but still retain disease standard scores for monomer secretion and NE inhibitory activity (**Fig. 7B**), highlighting critical SCV relationships that differentiate extracellular NE activity from intracellular polymerization.

Mapping the standard scores onto the structure (**Fig. 7C**) highlights the structural regions that have ‘healthy’ standard scores in terms of polymer formation but contribute to the deficiency in monomer secretion and NE inhibitory activity (**Fig. 7C**, arrows). Surprisingly, we previously showed that these structural areas are rich in variants that are only linked to lung-associated phenotypes and not liver-associated phenotypes in the patients^3^, suggesting that SCV precisely captures molecular mechanisms responsible for clinical phenotypes. Moreover, the residues that have disease standard scores below the mean for all the measured features (**Fig. 7D**, standard score<0) not only include the β-strand *s5A* and *RCL* that are critical for the anti-protease ‘loop-sheet’ insertion mechanism, but clearly illustrate the central role of β-sheet B formed by the long-range residue interactions of the β-strands contributed separately by N1 (*s6B*), M2 (*s2B-s3B*) and C3 (*s4B-s5B*), as a major determinant of liver and lung disease polymer, monomer and NE inhibitory phenotypes (**Fig. 7D**).

## Discussion

To evolve protein form and function requiring large and dynamic structural changes like the NE inhibitory activity of AAT, a delicate balance in sequence-to-function-to-structure relationships is needed to avoid protein misfolding and aggregation during nascent synthesis that leads to disease phenotypes. Using GPR, we have demonstrated that genetic variants in the population serve as fiduciary markers of evolutionary rules that can inform us on hidden biological mechanisms that drive the functional and structural design of the protein fold^1,3,41, 42^. The goal of understanding SCV relationships is not about the variant *per se*, but rather the use of natural variants as input to address mechanistically the multiple features, such as folding, secretion and NE activity in the case of AAT, that help us to understand sequence-to-function-to-structure relationships on a residue-by-residue basis across the WT protein sequence as to what can go wrong mechanistically in disease.

By profiling the SCV relationships of AAT variants, we capture the important role of the RCL and strand 5 in β-sheet A (*s5A*) that have been previously shown essential for protease inhibitor function in the lung and have been suggested to contribute to the classic ‘loop-sheet’ polymer generated using heat and chemically induced polymeric states *in vitro*^40, 59^. In addition, we now have discovered an underappreciated key element of AAT structural design, the β-sheet B that we posit serves as central feature for integration of all measured features including monomer secretion, intracellular polymer formation and NE inhibitory activity. Importantly, β-sheet B involves SCV relationships between β-strands derived from N-terminal (N1), middle region (M2) and C-terminal region (C3) of the polypeptide chain (**Fig. 7B**, red dotted circles)-suggesting that co- and post-translational long-range folding mechanisms differentially contribute to liver intracellular polymer disease and post-secretion loss-of-function lung disease. Consistent with this conclusion, the recent cryo-EM analysis of native Z-AAT intracellular polymer isolated from patient liver raised the possibility that, and different from the classic ‘loop-sheet’ polymer mechanism^40, 59–62^, the C-terminal two β-strands found in the β-sheet B (**Fig. 7B**, dotted red circles) insert into the β-sheet B of another AAT molecule to form the intracellular AAT polymer^57, 58^. Our SCV analysis based on evolved functional challenges to the fold found in the population supports this conclusion and further suggests that disruption of the β-sheet B stability through genetic variants in N1, M2 and C3 that impact the long range stability of this central integrating β-sheet core can also trigger intracellular polymer formation. Thus, an understanding of the short and long-range folding events captured through GPR based SCV relationships provide a quantitative mechanistic platform for developing novel approaches for management of maturation of AAT in disease.

As the secretory pathway involves flow through post-ER cellular environments, it is now clear that AAT has separately evolved unique sequence driven structural properties that manage its monomer secretion and extracellular function. Profiling these SCV relationships enabled us to differentiate the role(s) of residues facilitating nascent intracellular polymer formation from monomer secretion and NE inhibitory activity. For example, SCV relationships quantitatively demonstrate that a large sequence region in N1 does not contribute to the intracellular polymerization of AAT, rather to its rapid degradation - phenotypes associated with reduced or absence liver aggregate load but leading to typical late onset lung disease during aging^3^.

Interestingly, there is a glycosylation site on Asn 107 in this region, suggesting that the N-glycan dependent pathway may be uniquely poised for sensing functional-structural relationships in that region guiding its stability during nascent synthesis^11, 27, 28, 63, 64^. This conclusion is consistent with results that have demonstrated the importance of ERManI as the sole known genetic modifier in AATD^28^.

Numerous attempts have been made to prevent intracellular polymer in liver ER^37–39, 60–62, 65^. For example, small peptides have been developed to mimic the insertion of the cleaved RCL into β-sheet A to inhibit polymerization *in vitro*^60, 61^. However, given that NE inhibitory activity also requires a ‘loop-sheet’ insertion mechanism, these peptides block the NE inhibitory activity rendering questionable their use in the clinical setting^60, 61^. More recently, a small molecule GSK716^37–39^ binds at the top of *s5A* and prevents Z-AAT intracellular polymer formation, but it also blocks the NE inhibitory activity because it likely changes the orientation of the top end of *s5A* and impacts the ‘loop-sheet’ mechanism required for AAT activity. Therefore, a small molecule that can block intracellular polymer formation but does not impact (or even improves) NE inhibitory activity is a more attractive strategy for AATD clinical management. The discovery of a sequence regions contributing to intracellular polymer formation but not impacting NE activity empowered by GPR based SCV relationships may provide a promising target for drug development. These observations highlight the potential of GPR based SCV relationships to reveal hidden value dictating the highly complex, multi-dimensional properties involved in optimizing fold design for a particular functional feature using evolution as a guide and only a limited knowledge base of variation in the population responsible for disease.

From a more general perspective, we have previously demonstrated that SCV analysis can be used to understand the sequence-to-function-to-structure relationships for large membrane proteins with multiple domains such as CFTR^1,43^and NPC1^41, 42^ to decipher evolution-based design principles driving chloride channel function and cellular cholesterol management, respectively. Now, even for a small globular protein like AAT, SCV analysis of natural variants provides a rigorous sequence-to-function-to-structure interpretation at high resolution for the role(s) of different regions of the polypeptide sequence responsible for fitness. Thus, GPR analyses reveals that it is not the variant *per se*, but the weighted SCV relationships that are key to understanding principles of natural selection. Our results provide a rationale for how a linear genetic code can provide a highly dynamic template for testing possibilities essential for increasing fitness and biological diversity in response to the environment. Consistent with this interpretation we have recently^44^ suggested that biology is not so much about ‘quality control (QC)’ during nascent synthesis, but rather QC is only one facet of a broader ‘quality system (QS)’ where ‘quality assurance (QA)^44^ provides the key metric that dictates value in post-synthesis steps to assure evolution of diversity and optimal fitness. To understand complexity in fold design for function from an SCV perspective, we must approach the folding problem as has evolution- by embracing a computational framework such as GPR that allows us to see what nature sees to generate biology.

## Methods

### Reagents

DMEM, LHC-8 medium, F-12 medium, fetal bovine serum (FBS), penicillin streptomycin (P/S), and primocin TM were purchased from Invitrogen Life Technologies Corporation (Carlsbad, CA). FuGENE6 Transfection Reagent Kit was purchase from Promega (Madison, WI). Plasmid DNA purification kit were purchased from QIAGEN Inc (Valencia, CA). A goat anti-human AAT polyclonal antibody 80A was purchase from ICL Inc. (Anaheim, CA). The mouse anti-human monoclonal antibody 16f8 that preferentially recognizes AAT monomer was generated by the Scripps Research Antibody Development and Production Core. A mouse anti-human monoclonal AAT antibody 2C1 that preferentially recognizes AAT polymer were purchased from Hycult biotech (Wayne, PA). Recombinant AAT protein was purchased from Abcam (Waltham MA). Human neutrophil elastase (NE) was purchased from Innovative Research (Novi, MI). NE fluorescence assay substrate 2Rh110 (Z-Ala-Ala-Ala-Ala) was purchase from Cayman Chemical (Ann Arbor, MI). All other chemicals were purchased from Sigma Chemical (St. Louis, MO).

### Cell Culture

IB3 AAT null cells (generously provided by Dr. T. Flotte, University of Massachusetts Medical School, Worcester, MA) were cultured in LHC-8 medium containing 10% (v/v) fetal bovine serum (FBS) and 100 ug/ml penicillin streptomycin (P/S). Huh7.5 AAT knock-out (KO) cells (generously provided by Mark Brantly, University of Florida College of Medicine, Gainesville, FL.) were cultured in the DMEM/F-12 medium containing 10% FBS and primocin TM (100 ug/ml).

### AAT Variant DNA Constructs

AAT variant DNA constructs in a pcDNA3.1 (+) plasmid vector were generated by Quintara Biosciences (Cambridge, MA). All the plasmids were validated by independent sequencing (Genewiz, Inc (San Diego, CA)).

### Conformation specific AAT assays

IB3 null cells or Huh7.5 AAT KO cells were cultured in 96 well plates. AAT variant plasmids were transfected at 0.2 µg/well at a cell density of 2 x 10^4^/well. After 48 h transfection, cells were washed with phosphate-buffered saline (PBS) and incubated with 100 µl/well FBS free culture medium. After 3 h incubation, FBS free culture medium was collected. Cells were lysed in 80 µl buffer (25 mM Tris-HCl, pH 7.6, 150 mM NaCl, and 1% Triton) and harvested by centrifugation. 20 µl medium or cell lysates from each well was added to pre-coated and pre-blocked (BSA) 96-well plates containing goat anti-human AAT polyclonal antibody 80A. After an overnight incubation at 4^°^C, plates were washed 3X using PBST (phosphate-buffered saline/0.1% Tween). Conformation specific antibody 16f8 or 2C1 (mouse anti-human) was added to the plate and incubated for 2 h. After washing 3X with PBST, secondary HRP conjugated goat anti-mouse antibody was added and incubated 2 h. Following a 3X wash in PBST, TMB (3,3 ′,5,5 ′-tetramethylbenzidine) reagent was added into each well for 10 min and then stopped by 2M H_2_SO_4_. Plates were read by BioTek Synergy H1 Hybrid Reader (Santa Clara, CA) at 450 nm absorbance. The values were normalized by total protein levels in each well (measured by Bradford assay). Monomer levels of each AAT variant were normalized by WT monomer level. Polymer levels of AAT variant were normalized by Z-AAT polymer level. Commercially purchased AAT was serial diluted to provide an ELISA standard curve.

### NE inhibitory activity assay

Cell culture and medium collection were performed as described above in 96-well plate. 20 µl culture medium were added to pre-coated (goat anti-human AAT polyclonal antibody 80A) and pre-blocked (with BSA) plate. Following overnight incubation, medium was removed by washing 3X with PBST and plates incubated with NE at 5 ng/well for 2 h at 37^°^C. 25 pmol/well of 2RH110, a NE substrate, was added to plates and incubated for 1.5 h. Plates were read using a BioTek Synergy H1 Hybrid Reader (Santa Clara, CA) at excitation 485 nm and emission at 525 nm. The values were normalized by total protein levels in each well. The NE inhibitory activity of each AAT variant was normalized to WT AAT activity.

### Immunoblotting

Human AAT variants were transiently expressed in IB3 cell and/or Huh7.5 AAT knock-out cells. After 48 h transfection, cells were washed with PBS and then changed to FBS free culture medium for 3 h. Culture medium were harvested. Cells were washed twice with 1X with PBS and then lysed with 50 µl per well of cell lysis buffer (50 mM Tris-HCl, 150 mM NaCl, 1% (v/v) Triton X-100, a protease inhibitor cocktail (AEBSF, aprotinin, bestatin, E64, leupeptin, and pepstatin A each at 2 mg/ml) on ice for 30 min. Samples were collected and centrifuged at 20,000 x g at 4°C for 20 min, and supernatant collected. The protein concentrations of culture medium and cell lysate were determined by the Bradford assay (Bio-Rad, Hercules, CA). Culture medium or cell lysate samples were resuspended in SDS sample buffer containing β-mercaptoethanol and incubated at 95°C for 5 min. Samples containing 20 µg of total protein were separated using 10% SDS-PAGE, transferred to nitrocellulose, and immunoblotted with goat anti-human AAT antibody 80A. Detection was performed using chemiluminescence with horseradish peroxidase-conjugated secondary antibodies. Glyceraldehyde-3-phosphate dehydrogenase (GAPDH) was used as a loading control.

### Cycloheximide chase assay

Huh7.5 AAT knock-out cells were cultured and transfected with AAT variant DNA as described above. After 48 h transfection, cells were treated with cycloheximide (CHX) (50 µM) and chased for 0 h, 1 h, 2 h, and 4 h. Following the indicated chase period, cells were lysed in buffer (25 mM Tris-HCl, pH 7.6, 150 mM NaCl, and 1% Triton) and collected by centrifugation. Samples were quantitated for total protein levels by Bradford method and 15 µg total protein were separated using10% (v/v) SDS-PAGE, transferred to nitrocellulose, and immunoblotted with goat anti-human AAT antibody 80A.

### Variation spatial profiling (VSP) analysis

The VSP analysis was performed as previously described^1,41–43^ using gstat package (V2.0)^66, 67^ in R. VSP is built on Gaussian process regression (GPR) based machine learning. A special form of GPR machine learning that has been developed in geostatistics, Ordinary Kriging^68^, is used to model the spatial dependency as a variogram to interpolate the unmeasured value to construct the phenotype landscape for AAT. Briefly, AAT variants were positioned by their sequence positions in the polypeptide chain on the ‘*x*’ axis coordinate and their impact on a phenotype on the ‘*y*’ axis coordinate to the impact on another phenotype on the ‘*z*’ axis coordinate. Suppose the i^th^ (or j^th^) observation in a dataset consists of a value z_i_ (or z_j_) at coordinates x_i_ (or x_j_) and y_i_ (or y_j_). The distance h between the i^th^ and j^th^ observation is calculated by:

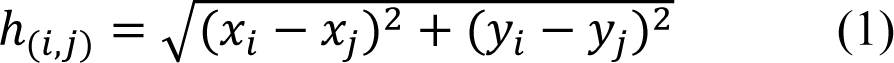

The *γ*(*h*)*-*variance for a given distance (*h*) is defined by:

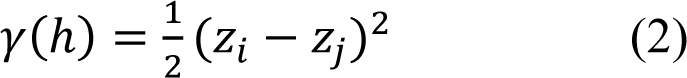

where (*h*)-variance is the semivariance (the degree of dissimilarity) of the *z* value between the two observations, which is also the whole variance of *z* value for one observation at the given separation distance *h,* referred to as spatial variance here. The distance (*h*) and spatial variance (*γ*(*h*)) for all the data pairs are generated by the equations (1) and (2). Then, the average values of spatial variance for each distance interval are calculated to plot the averaged spatial variance versus distance. The fitting of variograms were determined using GS+ Version 10 (Gamma Design Software) by both minimizing the residual sum of squares (RSS) and maximizing the leave-one-out cross-validation result (see below). The variogram enables us to compute the spatial covariance (SCV) matrices for any possible separation vector. The SCV at the distance (h) is calculated by C(*h*)= C(0) − *γ*(*h*), where C(0) is the covariance at zero distance representing the global variance of the data points under consideration (the plateau of the variogram). The approach aims to generate the prediction that has minimized estimation error (error variance) which is generated according to the expression:

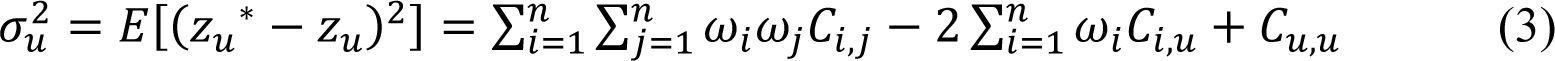

where *Z_u_^*^* is the prediction value while *z_u_* is the true but unknown value, *C_i,j_* and *C_i,u_* are SCV between data points i and j, and data points i and u, respectively, and *C_u,u_* is the SCV within location *u*. *ω_i_* is the weight for data point i. The SCV is obtained from the above molecular variogram analysis and the weight (*ω_i_*) solved from equation (3) is used for following prediction. To ensure an unbiased result, the sum of weight is set as one:

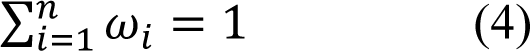

Equations (3) and (4) not only solved the set of weights associated with input observations, but also provide the minimized ‘molecular variance’ at location u which can be expressed as:

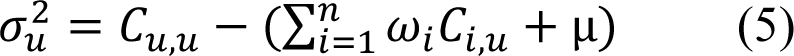

where *C_u,u_* is the SCV within location *u*, *ω_i_* is the weight for data point i, and *C_i,u_* are SCV between data points i and u. μ is the Lagrange Parameter that is used to convert the constrained minimization problem in Equation (5) into an unconstrained one. The resulting minimized molecular variance assessing the prediction uncertainty presents the confidence level of the prediction. With the solved weights, we can calculate the prediction of all unknown values to generate the complete fitness landscape by the equation:

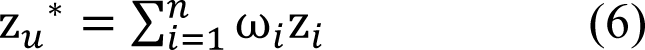

where z*_u_^*^* is the prediction value for the unknown data point *u*, *ω_i_* is the weight for the known data point, and *z_i_* is the measured value for data point i.

Leave-one-out cross-validation (LOOCV) is used to validate the computational model because of small sample size modeling^69^. In the LOOCV, we remove each data point, one at a time and use the rest of the data points to predict the missing value. We repeat the prediction for all data points and compare the prediction results to the measured value to generate the Pearson’s r-value and its associated p-value (ANOVA test performed in Originpro version 2020b (OriginLab)).

### Inverse variance weighting (IVW) analysis

Phenotype landscapes built based on a sparse collection of input variants map the full range of values describing function (based on the y- and z axis metrics) for the entire polypeptide sequence (x-axis). To obtain an averaged value of predicted phenotype for each residue, we use the reciprocal of GPR generated variance as weights to aggregate the phenotype values by using the following equation:

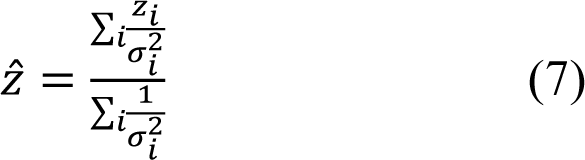

where *Z^* is the weighted mean value for each residue, *Z_i_* is predicted phenotype value at z-axis for every value on the y-axis, *σ^2^* is the GPR generated variance for each prediction. We repeat this process for all residues in the AAT sequence. The IVW averaged mean values for all the residues are mapped as a barcode based on color and to AAT structures (PDB:3NE4^70^ for AAT monomer; PDB:2D26^71^ for the AAT-NE complex and PDB:3T1P^57, 58^ for polymer). All atomic resolution structures were produced with the software of PyMOL.

### Standard score analysis

To compare the IVW averaged value on each residue for different phenotypes such as secreted monomer, intracellular polymer and NE inhibitory activity, we compute the standard score to normalize the phenotype value to the distribution of each phenotype for all the residues by using the following equation:

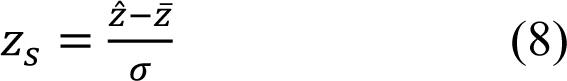

where *Z_S_* is the standard score for each phenotype of a residue, *Z^* is the IVW averaged value for each phenotype of that residue, *Z̅* is the mean of the distribution of the phenotype for all residues, and *σ* is the standard deviation of the distribution of the phenotype values for all residues. The standard score defines the number of standard deviations by which the phenotype feature is above or below the mean value of the phenotype feature for each residue in the AAT sequence plotted as line graph or as a standard score barcode as described in the Results.

## Acknowledgements

Grant support provided by NIH HL095524, DK051870, AG070209, AG049665, HL141810 and HG10881 to WEB. We thank Dr. Mark Brantly for providing the Huh7.5 AAT knock-out cells.

## Author contributions

All the authors contributed to the study design. C.W., P.Z., S.S. and X.W. developed the experimental assays. P.Z. and S.S. performed the experimental measurements for AAT variants. C.W. developed the computational methods. C.W. and P.Z. performed the computational analysis. C.W., P.Z., S.S., and W.E.B. wrote the manuscript.

## Supplementary Figure Legends

**Figure S1.**
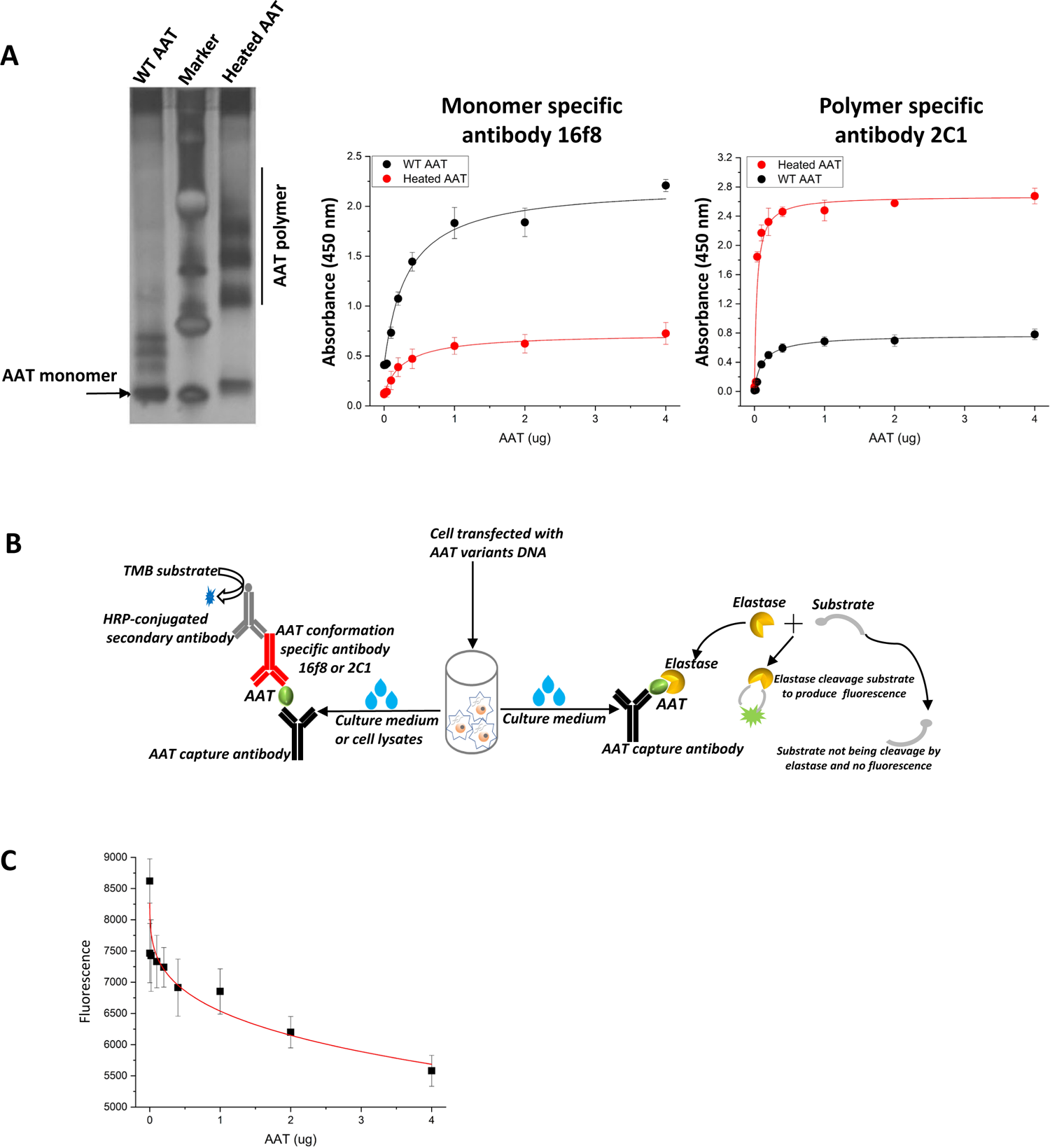
Conformation dependent antibodies and high-throughput assays used to measure different AAT phenotypes. (**A**) Shown is control WT monomeric and heat-treated polymeric AAT on native gel (left panel). Monoclonal antibody 16f8 generated shows strong interaction with WT monomeric AAT but not to heat-treated polymeric AAT (middle panel). Monoclonal antibody 2C1^13^ shows a strong interaction responding to the heated polymeric AAT but not WT monomeric AAT (right panel). (**B**) A schematic figure showing the high-throughput assays used to measure the intracellular and secreted monomer or intracellular and secreted polymer pools using conformational dependent antibodies (see **Methods**). The activity of secreted AAT is determined by using a fluorogenic substrate of NE (see **Methods**). (**C**) The fluorescence of the NE substrate is dependent on the protein level of AAT.

**Figure S2.**
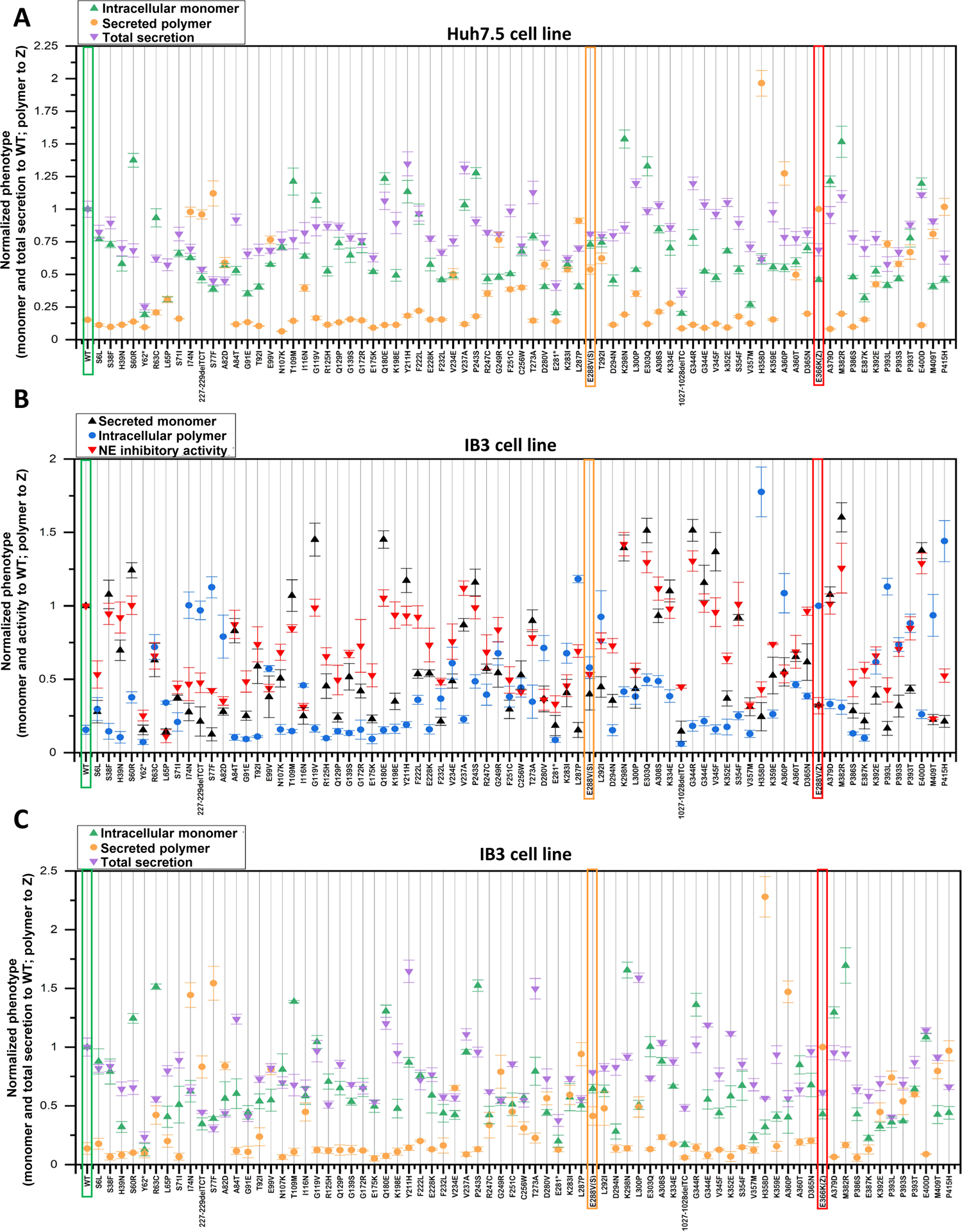
Phenotypes of AAT variants in different cell lines. (**A**) The intracellular monomer levels (blue), secreted polymer levels (orange) and total AAT protein (purple) secretion levels of 75 AAT variants transiently expressed in Huh7.5 cells (see **Methods**). (**B**) The secreted monomer levels, intracellular polymer levels and NE inhibitory activity for 75 AAT variants transiently expressed in IB3 cells. (**C**) The intracellular monomer levels, secreted polymer levels and total AAT protein secretion levels of AAT variants transiently expressed in IB3 cells. WT, E288V (S variant) and E366K (Z-variant) are highlight.

**Figure S3.**
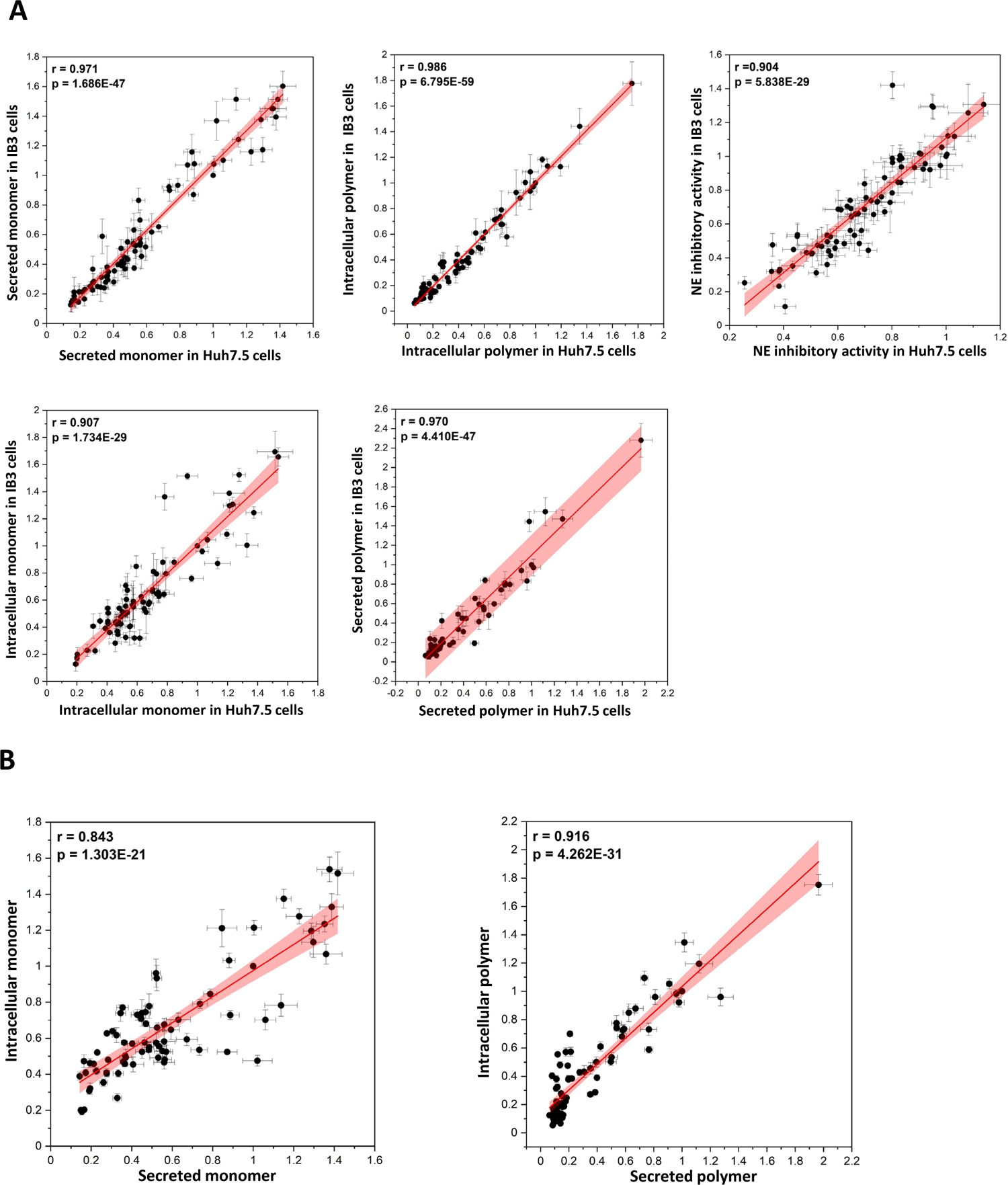
Phenotype correlations between different cell types. (**A**) The phenotype data of secreted monomer, intracellular polymer, NE inhibitory activity, intracellular monomer, secreted polymer for AAT variants are highly correlated between liver derived Huh7.5 cells and lung derived IB3 cells. Person’s r values and p values for each correlation are labeled. (**B**) In Huh7.5 cells, shown is the correlation between intracellular monomer levels with secreted monomer levels of AAT variants (left panel) and the correlation between intracellular polymer levels with secreted polymer levels for AAT variants (right panel). Pearson’s R values and p values for each correlation are labeled.

**Figure S4.**
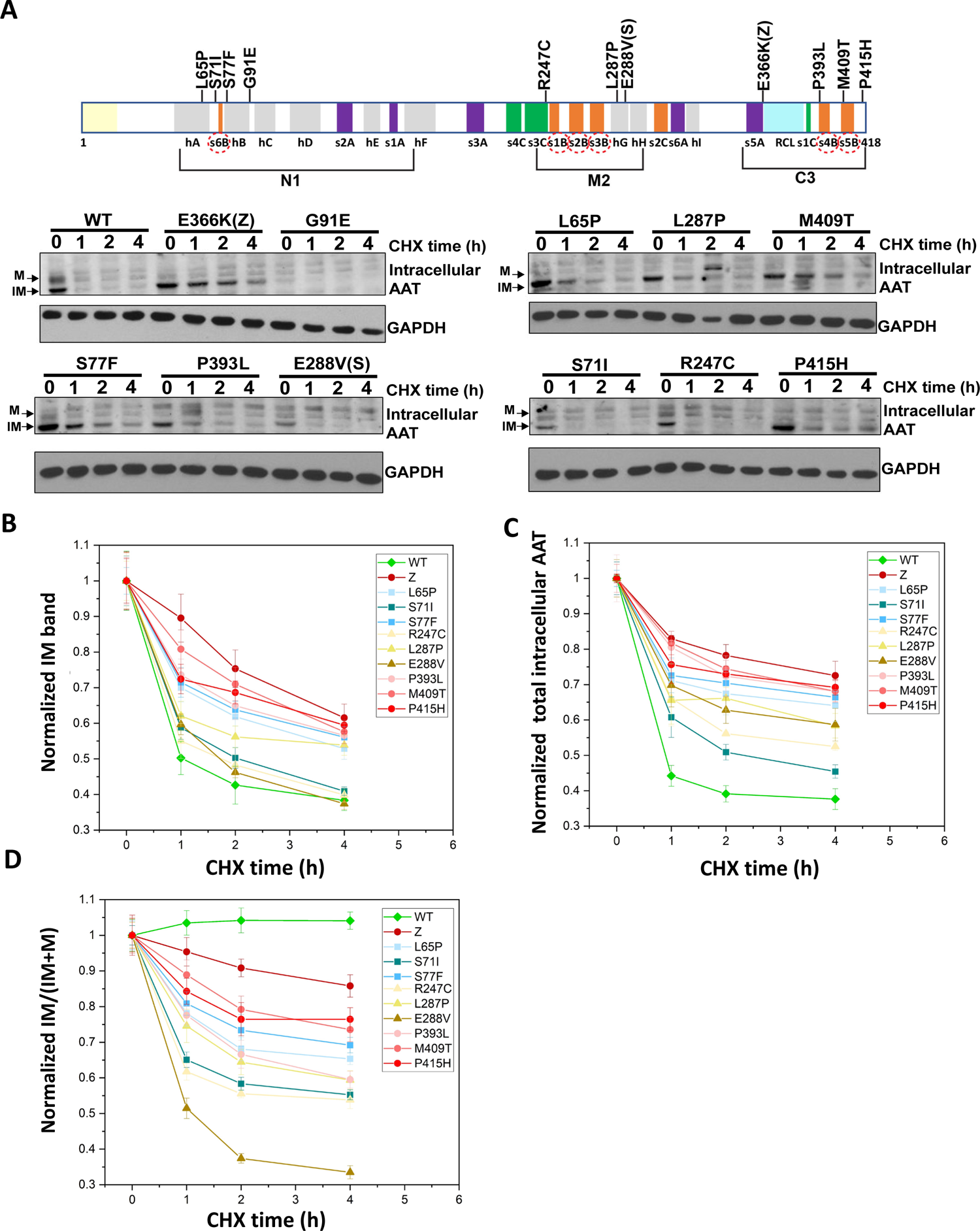
Cycloheximide (CHX) chase assay for N1, M2 and C3 variants using SDS-PAGE. **(A)** The indicated variants with secretion deficiencies found in N1, M2 and C3 regions were transfected in Huh7.5 AAT knock-out cells for 48 h and then treated with CHX (50 µM) for 0, 1, 2 or 4 h, respectively. Cell lysates were analyzed by SDS-PAGE and immunoblot (see **Methods**). <u>Im</u>mature bands (**IM**) representing the immature glycosylated AAT in the ER and the post-ER <u>m</u>ature (**M**) bands representing the post-ER mature glycosylated form of AAT are indicated. (**B**) The **IM** bands found at 1 h, 2 h and 4 h of CHX treatment were quantified and normalized to the value of the **IM** band at 0 h of CHX treatment for each variant to show rate of loss from the ER through either degradation or secretory pathways. G91E (panel **A**) is very unstable and degraded before CHX treatment and therefore not quantified. Variants in C3 region (red lines) show the slower rate of loss when compared with variants in N1 region (blue lines) and M2 region (yellow lines), suggesting that C3 variants are more stable inside ER when compared with other variants. (**C**) The total AAT protein ((**IM**) + mature band (**M**)) at 1 h, 2 h and 4h of CHX treatment was normalized to the levels of intracellular AAT protein found at 0 h of CHX treatment for each of the indicated variants. C3 variants (red lines) again show slower rate of loss for the total intracellular AAT when compared to N1 variants (blue lines) and M2 variants (yellow lines). (**D**) The ratio of quantitated density of gel bands (**IM**/(**IM+M**)) was determined for each of the indicated variants following CHX treatment for 1 h, 2 h, and 4 h, and normalized to 0 h of CHX treatment. C3 variants (red lines) show slower decrease on the rate of **IM**/(**IM+M**) over the CHX time course than N1 variants (blue lines) and M2 variants (yellow lines) indicate that the slower rate of loss for the ER fraction for C3 variants found in (**B**) is not because of secretion, but because C3 variants form more stable conformations that are harder to be degraded, highlighting the different structural stabilities of the folding species impacted by variants in different regions like N1, M2 and C3 along the AAT polypeptide sequence.

